# Honest signalling in predator-prey interactions: testing the resource allocation hypothesis

**DOI:** 10.1101/2024.12.06.627197

**Authors:** Emily Burdfield-Steel, Cristina Ottocento, Miriam Furlanetto, Bibiana Rojas, Ossi Nokelainen, Johanna Mappes

**Affiliations:** Institute for Biodiversity and Ecosystem Dynamics, University of Amsterdam, Amsterdam, The Netherlands; Organismal and Evolutionary Biology Research Program, University of Helsinki, Helsinki, Finland; Department of Biological and Environmental Science, University of Jyväskylä, Jyväskylä, Finland; Department of Interdisciplinary Life Sciences, Konrad Lorenz Institute of Ethology, University of Veterinary Medicine Vienna, Vienna, Austria; Open Science Centre, University of Jyväskylä, Jyväskylä, Finland

**Keywords:** aposematism, chemical defence, life-history, resource allocation, predator-prey interactions, Lepidoptera, wood tiger moth, pyrazine, warning signal, melanin, honesty signalling

## Abstract

Warning signals are honest if they reliably deliver information about prey unprofitability to predators. One potential mechanism that may create and maintain a positive relationship between the strength of signals and defence is the resource allocation between these costly traits. Here, we test this hypothesis using the wood tiger moth *Arctia plantagini*s, whose females’ red hindwings are a warning signal to predators but show considerable variation in colouration within populations. These moths also produce a defensive chemical that is known to influence avian predator attack risk. Using dietary manipulations, image and chemical analyses, and experiments with ecologically relevant predators we demonstrate that protein availability during development can influence the strength of both the primary warning signal and the secondary defence. Our results show that females raised on a high-protein or *ad libitum* natural diet produced more distasteful defensive fluids than those raised on a low-protein diet or subjected to periodic food deprivation. While the patterning of the warning signal was unaffected by food deprivation, its efficacy was diminished in moths raised on a low-protein diet. However, this change was imperceptible to avian predators. Critically, resource availability influenced the relationship between signal strength and defence: moths on a high-protein diet displayed a positive correlation between warning signal strength and unpalatability, whereas this correlation was absent in moths raised on a natural diet. These findings show that resource availability can weaken the reliability of warning signals as an indicator of an individual’s defensive capabilities, highlighting the complex interplay between ecological conditions and the evolution of honest signalling.

## Introduction

The stabilising selection on prey warning signals imposed by predation is a central pillar of warning signal theory (Sheppard et al., 1985; Fisher, 1930). Given that most organisms benefit from deterring a potential predator before it attacks, many prey species advertise the defences they possess with distinctive warning signals (Poulton, 1890; Rojas et al., 2015). These warning signals then become associated with the unprofitability of the prey for the predator, either by associative learning within a generation or across generations via evolved aversions to certain stimuli (Rojas et al., 2015; Ruxton et al., 2019). To facilitate predator learning, warning signals are expected to be easy to detect and remember, conspicuous and, above all, consistent (Speed, 2000). For instance, predators have been shown to learn to avoid certain signals faster when they encounter them more often, creating selection against rare signals (Lindström et al., 2001; Gordon et al., 2021, although this may be dependent on the total number of prey available, see Rowland et al., 2010). As a result, predators are predicted to exert stabilising selection on aposematic signals in local prey populations, removing those that deviate from the norm (Mallet & Barton, 1989). Largely, this prediction has proven true. There are numerous examples of mimicry rings in which multiple species, sometimes from highly diverged taxa, share similar warning signals – such as red-and-black or yellow-and-black colouration – to share the costs and benefits of predator education (Wilson et al., 2012; Twomey et al. 2013; Garg et al., 2019; Nokelainen et al., 2023). However, there are also several examples of warning signal polymorphism within populations, and variation in signal strength or consistency is widespread (Briolat et al., 2019). There are many potential explanations for such variation. One mechanism that has received increasing interest is when resource allocation decisions lead to the honest signalling of prey defence and hence prey unprofitability (Blount et al., 2009; Summers et al., 2015).

Honest signalling in aposematism arises if there is a positive correlation between a primary signal (such as colouration) and a secondary defence (such as toxicity) (Summers et al., 2015). While *qualitative* honesty, a link between possessing both the signal and the defence, forms the very basis of aposematism, *quantitative* honesty, where the strength of the signal and defence are positively correlated, is more controversial. Some species do show positive correlations, including ladybirds (Bezzerides et al., 2007; Blount et al., 2012), paper wasps (Vidal-Cordero et al., 2012) and poison frogs (Maan and Cummings, 2012); others show negative correlations (Wang, 2011) or none (Briolat et al., 2018; Stuckert, Saporito and Summers, 2018; Briolat et al., 2019). Nevertheless, a recent meta-analysis concluded that there was a positive correlation between warning colour and chemical defences, and that this held across all scales, from within-population to across species (*n* =22 different species; White and Umbers, 2021). Despite this, the mechanisms responsible for promoting and maintaining honest signalling in aposematic species remain unclear. A so-called handicap model of aposematism suggests that honesty requires differential costs of signalling, that is, that strong signals must be more costly to produce than weak ones (Yachi, 1995). However, “costs” can take many forms. For example, at least three different mechanisms that involve signal costs have been proposed to facilitate honest warning signals in prey (Holen and Svennungsen, 2012): resource allocation trade-offs, go-slow behaviour in predators, and costs of detection alone. The detection cost hypothesis (when the probability that the predator spots the prey is enough to maintain the honest signalling equilibrium) was shown to be limited (Holen and Svennungsen, 2012). Go-slow predation, and the closely related taste-rejection (Guilford, 1994), have been suggested to be key mechanisms for punishing cheaters (i.e., lower-defended individuals) and have received some experimental support (Skelhorn and Rowe, 2006). These mechanisms do not, however, explain how the variation initially arises.

Resource allocation, in contrast, provides a potential mechanism for honest signalling, proposing that both signal and defence may draw on a shared pool of resources, creating trade-offs (Blount et al., 2009; Holen and Svennungsen, 2012). A previous study on ladybirds showed evidence that the availability of resources influences both chemical defences and their colouration, as well as a positive correlation between the two when resources are limited (Blount et al., 2012). Additionally, in females (the larger and hence perhaps more resource-limited sex), the correlation between the carotenoid concentrations and total levels of the alkaloid coccinelline was positive, but that was not the case in males (Blount et al., 2012). However, said study did not measure the response of predators to these alkaloid levels directly, which is one additional complication in the study of honest signalling: determining if observed variations in both colour and defence are detectable, and usable, by predators. The crucial condition for warning signal honesty is that *both* signal and defence traits are under selection by predators. Thus, while the existence of positive correlations between colour and chemical defences is highly suggestive of honest signalling (e.g., White and Umbers 2021), both must also be shown to influence predator behaviour to confirm it. Many studies still use proxies such as bioassays, and even direct measurements of the abundance of certain chemicals, to determine the strength of chemical defences (White and Umbers, 2021). However, neither of these may give an accurate view as it is not always known if predators’ response to prey defence is linear (Lawrence et al. 2019) and whether the within population variation in defence is detected by predators (Ottocento et al., 2022) and hence under selection. Therefore, studies that include accurate information on the responses of ecologically, and evolutionarily, relevant predators to both the colouration and chemical defences of the studied species are vital to further our understanding of honest signalling.

The wood tiger moth *Arctia plantaginis* is a well-suited system for such studies, as both its colour and chemical defences have been extensively investigated. Birds have been shown to vary their attack behaviour in response to variation in the moths’ hindwing colouration (Lindstedt et al., 2011; Nokelainen et al., 2014; Rönkä et al., 2018a; Rönkä et al., 2020), and their chemical defences have been characterised (Rojas et al., 2017; Burdfield-Steel et al., 2018) and can be tested in isolation on ecologically relevant predators (Burdfield-Steel et al., 2019; Rojas, Mappes and Burdfield-Steel, 2019; Winters et al. 2021). The species is also a good candidate for resource allocation trade-offs as it is polyphagous, meaning that resource quality can vary greatly depending on the host plant, and a capital breeder, so additional resources are not acquired in the adult life-stage. Proteins are crucial components of the diet of herbivores such as *A. plantaginis*: they are necessary precursors to the production of the pigments that form the warning colouration of the wood tiger moth (Lindstedt, Morehouse, et al., 2010a; Lindstedt, Suisto and Mappes, 2020) and used in the synthesis of chemical defences in insects (Morgan, 2004). Resource availability during development has already been found to influence the strength of wood tiger moth warning colouration (Lindstedt, Suisto and Mappes, 2020) and, in a separate experiment, food deprivation was shown to reduce the strength of their chemical defences confirming its costly nature (Burdfield-Steel et al., 2019). However, how resource allocation affects both the warning signal, and the chemical defences simultaneously has not been rigorously tested.

Here, we tested the effect of resource availability during development on both the warning colour (primary defence) and chemical defence (secondary defence) in the aposematic wood tiger moth. We used four dietary manipulations: natural diet (*ad libitum Taraxacum spp.)*, food deprivation (*Taraxacum spp*. but food-deprived for one day), high-protein content, and low-protein content. We used multispectral image analysis and bird vision model analysis to look for colour changes in the hindwings, which are the primary visual warning signal in this species, and tested their defensive fluids, in the absence of colour cues, on ecologically relevant predators to confirm that changes in the chemical composition of the fluids impacted predator deterrence.

We hypothesised that the female aposematic signal in this species is *quantitatively* honest: when both traits draw on the same pool of resources, individuals that have fewer resources during development will show both weaker visual signals and weaker chemical defences. Thus, we expect that individuals raised on high-protein conditions will have (1) a conspicuous warning signal with redder hue, redder colour detectable by avian predators, higher saturation, lower brightness (orange hindwings are brighter than red hindwings but are less conspicuous to predators; Lindstedt et al., 2011), and higher melanin amount; (2) better chemical defences (higher pyrazines levels, with less variability, and increased aversion induced in birds); and (3) shorter developmental time, larger wings and heavier pupal mass compared to the individuals raised on the low-protein diet. Additionally, we predicted the same pattern when comparing moths raised on a natural *ad libitum Taraxacum spp.* diet to individuals raised in periodic food deprivation (4 and 5). Finally, aposematic signals are predicted to be honest when resources are limited, leading to a positive correlation between resource supply and resource allocation (Blount et al., 2009). Within treatments, we hypothesised that (6) individuals raised on the low-protein and natural food-deprived diets, but not those raised on high-protein and *ad libitum* natural diets, would show a positive correlation between chemical defences defences and warning colouration, as the trade-offs become more evident in the resource limitation environments and thus, expected to “produce signal honesty”.

## Methods

### Study species

*Arctia plantaginis* (formerly *Parasemia plantaginis*; Rönkä et al., 2016) is a diurnal moth, widespread across the northern hemisphere. Male hindwing coloration varies discretely within and between populations, and can be yellow, white, black, or red. Hindwing colouration in females, by contrast, varies continuously from yellow to red in most populations throughout their range. This species produces two types of defensive fluid when attacked, one from the thorax, which is targeted to avian predators, and one from the abdominal tract, which is targeted to invertebrate predators (Rojas et al., 2017). The thoracic defensive fluid contains methoxypyrazines, SBMP (2-sec-butyl-3-methoxypyrazine) and IBMP (2-iso-butyl-3-methoxypyrazine), which are produced *de novo* (Burdfield-Steel et al., 2018). As wood tiger moths are capital breeders they do not feed as adults and must acquire all their resources during the larval stage. While male colouration is genetically determined (Brien et al., 2022), female colouration shows environmentally induced variation (Lindstedt et al. 2010a) despite being highly heritable (Koch et al., 2024; Lindstedt et al., 2016). Crucially, there is already evidence that early-life resource availability can influence both female colouration and chemical defence (Burdfield-Steel et al., 2019; Lindstedt, Suisto and Mappes, 2020), and that this variation is detectable by predators (Lindstedt et al., 2011). Thus, this species represents an ideal system in which to test the effects of resource availability on strategic allocation and signal honesty.

### Diet manipulation

Adult moths were taken from a stock population of moths, founded in 2011 from wild individuals collected in Estonia and maintained at the University of Jyväskylä. To create families that covered the range of colours seen in wild females, males and females were paired up based on female colour, and the colour of the male’s sisters. Female hindwings, which exhibit continuous colour variation from yellow through to red, were attributed to a numerical classification by eye, ranging from 1 to 5 (yellow to red, see Nokelainen et al., 2022). This classification is consistent with the range of variation perceived by avian predators (Lindstedt et al., 2011). After mating occurred, the males were removed, and the females were allowed to lay eggs. Upon hatching, larvae of the same family were kept together for the first 14 days and fed with a mixed diet of dandelion (*Taraxacum spp*.) and lettuce leaves. Larvae are polyphagous, but *Taraxacum spp.* has been found to be amongst the most common constituents of their diet. Every family was then counted and split into four different treatments. One fed *ad libitum* with *Taraxacum spp*., (hereafter referred to as *ad libitum* natural diet), one fed with *Taraxacum spp.* but food deprived for 24 hours once per week, one fed on a low-protein artificial diet, and one fed on a high-protein artificial diet (see supplementary material, Table S1). Depending on the initial number of eggs hatched per each family, a total of 10, 20 or 40 individuals were assigned to each treatment. In total, the experiment involved 8 families for treatments with natural diet (180 individuals per treatment) and 10 families for artificial diet treatments (240 individuals per treatment). However, as not every family was large enough to be split in four treatments, three families (5, 6, and 11) were kept only in artificial diet treatments and one family (7) was reared only in the natural diet treatments. Larvae were reared in boxes of 10 individuals until they started pupating; at that point, they were separated and kept in individual containers to reach adulthood. The experiment took place in a greenhouse at the University of Jyväskylä in Central Finland (62°N, 26°E) from May to August 2017. The temperature in the greenhouse varied between 20°C and 30°C, decreasing to 15°C-20°C during the night; daylight lasted for approximately 20 hours. Every box was checked and watered daily, cleaned when needed; fresh food was added *ad libitum* and old leaves were removed. Pupae were kept at room temperature (25°C ± 2°C) and their condition was checked daily. The individuals were weighed when they reached pupation, as this is a good proxy for adult body weight (Ojala et al., 2005). Hatching, pupation and adult emergence dates were recorded for each individual. Uncommon behaviours such as larval cannibalism and early mortality were also monitored. Life-history traits (developmental time per days, pupal weight, survival, hind- and fore-wings size), and colour measurements were recorded for every individual. Additional measurements of the strength of chemical defences and colouration were taken for at least three adult females per family per treatment (see below). Males emerging from the dietary treatments were also sampled but, as the focus of this study was female defence and colouration, we include only data on females. The analyses were performed on a dataset including females that were tested for both colour, melanin and chemical defences, except for survival rate differences, which were calculated on the entire dataset of individuals reared in the experiment, as sex could not be determined if an individual died prior to eclosure. Predator assays were conducted with fluids from females emerging from the same experiment; however, as the very small volume of fluids from a single female can only be used for one purpose, we do not have pyrazine measures for these individuals and vice versa.

### Colour measurements

Moths were kept at 7°C for 1-14 days after emerging as adults. For this study 149 females were sampled for their chemical defences (101 individuals were used to measure the amount of pyrazine, and 48 individuals - 11 from the food-deprived diet; 10 from the *ad libitum* natural diet; 12 from the low-protein diet; 15 from the high-protein diet - were used to test the correlation between the warning signal and the efficacy of the chemical defences against birds), then set for photography using needles on wooden boards. After drying, set specimens were photographed in full spectrum light under controlled conditions. A light source (Exo-Terra SunRay 75W), emitting in the full visible light spectrum, was used to illuminate the individuals during the photography. Photographs were taken in RAW format with a Samsung NX1000 digital camera, customised into full spectrum range and equipped with Nikon EL-Nikkor 80 mm enlarging lens, using an established protocol from Nokelainen et al., 2017. All the image analysis were performed with the Image calibration and analysis toolbox plugin (Troscianko and Stevens, 2015) for ImageJ (v. 1.50f), creating multispectral images that combined photographs taken in the UV and visible spectra. From every image, a set of regions of interest was selected: (a) whole forewing area (b) forewing melanised region (c) whole hindwing area (d) hindwing melanised region and (e) hindwing coloured region. These were reported in a dataset using the batch analysis tool (ImageJ), selecting the Visible R luminance channel and running it through no visual system, in order to observe the raw camera responses as objective reflectance data. The RGB values were converted into hsv (hue-saturation-value) colour space using the rgb2hsv in R. As hue is a circular value, we added an offset to all hue values to make the red degrees comparable. The relative proportion of the melanised area was calculated as follows for both fore- and hindwings: [(wing melanised region area in pixels) / (whole wing area in pixels)].

To assess how avian predators are expected to perceive the hindwing colouration of *A. plantagini*s, we analysed the ultraviolet (U) spectra along with blue (B), green (G), red (R) channels, and luminance (Lum) of the wings. The same individuals and images were processed for this analysis, using the blue tit, *Cyanistes caeruleus,* visual system D65 daylight, and 300-700 nm model in ImageJ (Troscianko and Stevens, 2015).

### Quantification of chemical defences

After individuals from the four dietary treatments emerged from pupae, they were given a drop of water to hydrate and put at 4°C to slow the metabolic rate down. To collect the thoracic fluid, the moths were kept at room temperature (20°-25°C) for at least 30 minutes. The thoracic fluid was extracted by gently squeezing the moths below the head, collected using capillaries of 10 μm in diameter and stored at -18°C in glass tubes until used for either gas chromatography – mass spectrometry (GC-MS) analysis, or trials with live birds. The volume of fluid collected from each adult was also recorded. The pyrazine content of the fluids was tested using GC-MS following the protocol previously used by Rojas et al., (2017) and Burdfield et al., (2018).

### Predator assays

We used 64 blue tits (*Cyanistes caeruleus*) as a model predator as they are common in Finland, easy to capture and keep in captivity for short periods of time. As an omnivorous and insectivorous bird, this species is a natural predator of the wood tiger moth and shares a similar range of distribution in Europe. Finally, it has been used in several previous studies of wood tiger moth predator defence (Nokelainen et al., 2012; Rojas et al., 2017; Rönkä et al., 2018b; Burdfield-Steel et al., 2019). The birds used in this experiment were caught at Konnevesi Research Station, in Central Finland, using baited traps and maintained individually in plywood cages with a perch, water bowl and food *ad libitum*, and kept on a 12:12 h light:dark cycle. Each bird was weighed before and after the experiment, ringed, and its sex and age were determined before being released to the same place of capture. Birds were used with permission from the Central Finland Centre for Economic Development, Transport and Environment, and licensed from the National Animal Experiment Board (ESAVI/9114/04.10.07/2014) and the Central Finland Regional Environment Centre (VARELY/294/2015). All experimental birds were used according to the ASAB/ABS Guidelines for the treatment of animals in behavioural research and teaching. All birds were introduced to eating bait (oat flakes) while still in their home boxes. They were then familiarised with the experimental box and trained to eat the bait from the feeding hatch prior to the experiment. Only when birds were observed eating the baits in the boxes were the trials started. Each bird was used only for one assay which consisted of four trials, one after another with 5-minute intervals between trials. The first and last were control trials in which the bait was soaked in water; this was done to establish that the birds were motivated to feed. In the second and third trials birds were exposed to the chemical defences of a single moth. We rehydrated samples with 15μl of water. In all trials 7μl of fluid (either water or rehydrated defensive fluid) was pipetted onto the bait and allowed to soak in for 5 minutes before being presented to the bird. The maximum duration of each trial was 5 minutes, and the trial ended two minutes after the bird had eaten the whole bait. If the bird did not eat the bait in the first trial within 5 minutes, the test did not continue. The final trial ensured that the birds were still motivated and hungry, and thus any reaction to the middle trials was due to the fluid sample rather than satiation. For each trial, the predators’ response was measured. As a proxy for distaste, we recorded the proportion of the bait eaten, the number of times the birds cleaned their beak (recorded as the number of bouts of beak cleaning, i.e., against any surface), the latency to approach (the time that took for the bird to get close to the plate with the bait), the latency to eat after approaching (the time between approach and attacking the time) and the latency to eat (the time from when the bird sees the bait on the white plate to the time from when the bird starts to eat the bait). All trials were recorded with a camera positioned on the roof of the box, allowing any missed behaviours to be checked after the trial was finished. Birds were only used for a single assay and had not been previously used in any other experiment involving exposure to the moths’ defensive fluids.

### Female fecundity

In a follow-up experiment the high- and low-protein diets were repeated in the subsequent summer with Estonian stock families, and mating success, egg number, and larval hatching success were assessed (see supplementary material, paragraph 7, for details).

#### Statistical Analysis

The statistical analyses were done with the software R v. 4.1.2 (R Core Team, 2022) using the RStudio v. 1.2.1335 interface (RStudio Team, 2019) and R 4.1.0. We set the level of significance in all analyses at *p < .05*.

We used planned contrasts (package *car*; Fox et al., 2022) across all measures to compare the different diet treatments. We compared (1) natural diet *ad libitum* (DH) and natural diet food-deprived (DL) to measure the effect of food deprivation; (2) high (PH) and low (PL) protein diets, to examine the effects of protein availability; and finally (3) both natural diets with both artificial diets, to compare the two different feeding regimes.

### Life history

Life-history measures were only analysed for females. All measures were analysed with treatment as a fixed factor and family as a random factor to account for any family-level effects. Development time was analysed with a mixed effects Cox model, with a Gaussian distribution. Pupal weight was analysed as a general linear mixed model with link=log distribution, using the *lme4* package (Bates et al. 2015). The correlation between development time and pupal weight was tested with Spearman’s rank correlations (Ojala et al., 2005). Survival was modelled as a general linear mixed model with a binomial distribution (package *lme4*; Bates et al. 2015).

### Colour measurements

Hue, saturation, brightness, wing size and the proportion of the wing melanised, were analysed with linear mixed models with treatment as a fixed factor and family as a random factor, using the *lme4* package (Bates et al. 2015). The same package and modelling approach were applied to analyse the UV, blue, green, red channels and luminance of the avian vision model, with treatment as a fixed factor and family as a random factor.

### Chemical analysis

The abundance of SBMP and IBMP in the moth’s defensive fluids was analysed with linear mixed models (package *lme4*; Bates et al. 2015), with treatment and volume of fluid as fixed factors and family as random factor. Bartlett test (Bartlett, 1937) was applied to compare variance in the amount of each pyrazine among treatments. Spearman’s rank correlations were used to test how wing colouration (hue, saturation and brightness), wing size, and amount of melanin in the wings correlate to amounts (ng) of SBMP and IBMP in the wood tiger moth’s chemical defences.

### Predator assays

Predator responses to the chemical defences of the moths were first compared to a control (i.e., water) to determine whether the chemical fluids from the different treatments elicited an aversive reaction. To test the proportion of bait eaten we used a beta distribution: transforming first the proportion to be between 0 and 1 and then using package glmmTMB (Brooks et al., 2017) with family =beta_family(link= “logit) for beta regression models with random effects. We used Generalized Linear Mixed-Effects Models (glmer; package lme4, Bates et al. 2015) fit by maximum likelihood (Laplace Approximation) and family negative binomial with trial duration set as offset to test the effect of the moths’ chemical defences on beak wiping behaviour. To test the latency to approach, the latency to eat after approaching and the latency to eat, we used the Cox mixed-effects model using the package *coxme* (Therneau, 2020), fit by maximum likelihood. The predators’ behaviours were set as response variables; the different types of diet treatments were set as fixed factors, and bird ID was set as a random factor. The treatments were then compared with the planned contrast excluding the water control group. We used Spearman’s rank correlations to investigate whether the wing colouration (hue, saturation and brightness), size and amount of melanin correlate with predator responses (proportion of bait eaten, beak wiping, latency to approach, latency to eat after approaching, latency to eat) on the chemical defence. Additionally, we tested Spearman’s rank correlations between predator responses and the UV, red, green, blue channels and luminance from the avian vision model.

## Results

### Diet manipulation-life-history effects

A total of 840 larvae were followed from eggs to adulthood or death. Of these, 757 reached the pupal stage and 698 emerged as adults, of which 332 were females and subsequently included in the life history analysis. Diet restriction strongly influenced developmental time from larva to pupation (Figure 1AS, Supplementary Table 2S). Larvae fed on an *ad libitum* natural diet, took less time to develop and reach pupation than those fed on a food-deprived natural diet (z= -3.65, *p<0.001*), as did larvae fed on the low-protein artificial diet when compared to the high-protein artificial diet (z= -3.40, *p<0.001*). Moreover, larvae fed on both artificial diets took longer to reach pupation than those fed on natural diets (z= -13.48, *p<0.001*). Pupal weight also showed clear differences between the natural and artificial diets (Supplementary Figure 1BS; Supplementary Table 3S), as artificial diets produced heavier pupae than natural diets (t= 9.39, *p<0.001*). There was also a significant difference between the *ad libitum* and food-deprived treatments fed on a natural diet (t= -2.08, *p= 0.04*), as the food-deprived larvae became slightly smaller pupae. However, there was no significant difference between the high and low-protein diets (t= 0.35, *p= 0.73*). The relationship between development time and pupal weight also differed between the treatments (Supplementary Figure 1CS). The artificial diet treatments showed positive correlations between development time and pupal weight (Spearman’s rank correlation of 0.66 and 0.62 for the high and low-protein treatments respectively), but the natural diet treatments did not (Spearman’s rank correlation of -0.06 and -0.21 for *ad libitum* natural diet and food-deprived respectively). Overall, there was no difference in survival to adulthood between the natural diet- and artificial diet fed moths (z= 0.23, *p= 0.82*), or between the moths fed on *ad libitum* natural diet versus those that were food-deprived (z= 0.31, *p= 0.76*). However, the survival rate was significantly lower in the low-protein artificial diet than in the high-protein diet (z= -3.69, *p<0.001*). This was likely a result of an increased level of sibling cannibalism observed in the artificial diet treatments (Furlanetto pers. obs.), and the number of individuals from the low-protein treatment that failed to pupate.

**Figure 1.**
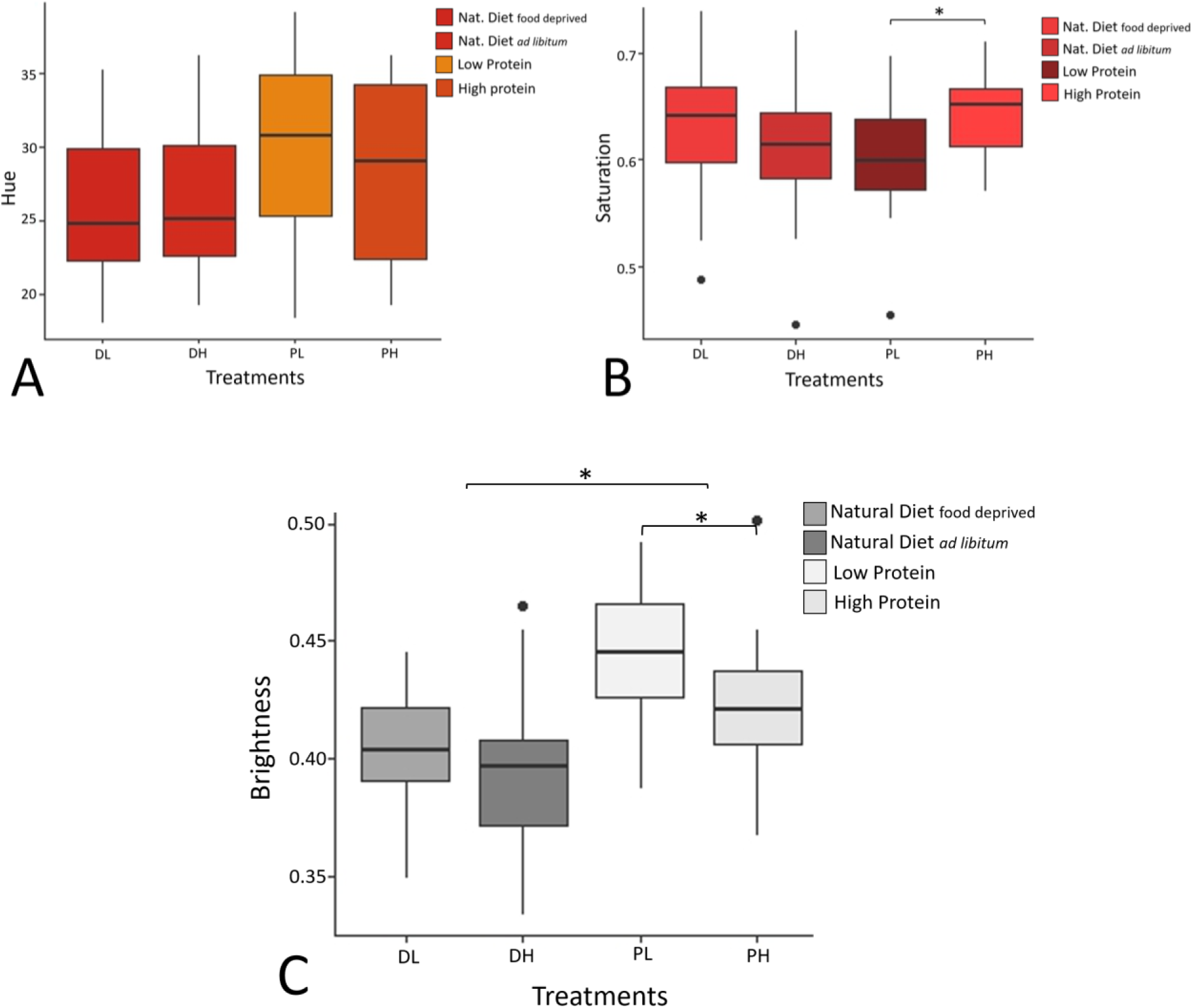
**A)** Hindwing hue, **1 B)** Hindwing saturation and **1 C)** Hindwing brightness of female moths from the four different diets. The colours of the boxplots denote the differences in colour metrics in the female hindwings among treatments. Boxes show the median and the 25th and 75th percentiles of data distribution. Vertical lines show the data range. (DL = natural diet food-deprived, DH = natural diet *ad libitum*, PL = Low-protein artificial diet, PH = High-protein artificial diet)

### Colour measurements

When looking at absolute colour, hue values did not differ significantly between individuals raised on different treatments (*p > 0.05* for all contrasts; see Table 4S, supplementary material; Figure 1A). Saturation (intensity of colour) showed significant differences between moths reared on high and low-protein diets - with moths reared on high-protein diet having more saturated hindwing colouration than those reared on the low-protein diet (t = 3.10, *p = 0.003;* Figure 1B). Saturation did not differ, however, between females from *ad libitum* and food-deprived natural diets (t = 1.06, *p = 0.29;* Figure 1B) nor between females from the natural and artificial diets (t = 0.28, *p = 0.78;* Figure 1B). Moths raised on the high-protein diet showed lower hindwing brightness compared to those on the low-protein diet (t = -3.00, *p = 0.004;* Figure 1C), but no such difference was found between moths from *ad libitum* and food-deprived diets (t = -1.08, *p = 0.28;* Figure 1C). However, moths raised in the artificial protein diets were brighter than moths raised in the natural diets (t = 5.55, *p < 0.001*; Figure 1C). The amount of melanisation in the hindwings was higher in moths reared in the artificial diets than in moths raised in the natural diets (t = 3.00, *p = 0.0035;* see Table 7S, supplementary material). Similarly, the amount of melanisation in the forewings was greater in females from the artificial protein diets than females from the natural diets (t = 2.66, *p = 0.0094;* see Table 8S, supplementary material). There were no differences in the size of the hindwings between treatments (*p > 0.05* for all contrasts; see Table 9S, supplementary material). There was however a difference in the size of the forewings between the natural and artificial diets, with moths reared on artificial diets having larger wings than moths reared in natural diets (t = 2.20, *p = 0.03;* see Table 10S, supplementary material).

When looking at the bird vision model, UV values did not differ between individuals raised on different treatments (*p > 0.05* for all contrasts; see Table 11S, supplementary material). Similarly, there were no significant differences in the blue colour channel values across treatments (*p > 0.05* for all contrasts; see Table 12S, supplementary material). Individuals raised on the artificial protein treatments showed a tendency to present higher green values than those reared in the natural diet groups (t = 2.006, *p = 0.051*; see Table 13S). No significant differences in the green channel were detected between other treatments. Although not significant, moths from the artificial protein diets present slightly higher red channel values compared to moths raised in the natural diets (t = 1.891, *p = 0.065*; see Table 14S). No significant differences were found in the other treatments. Luminance values also showed no significant differences between treatments (*p > 0.05* for all contrasts; see Table 15S, supplementary material).

### Quantification of chemical defences

Overall, moths reared on the artificial diets produced higher amounts of both pyrazines than those fed on natural diets (SBMP: z= 2.443, *p= 0.0146* and IBMP: z= 1.992, *p= 0.046*). No significant differences in the abundance of SMBP or IBMP were found either between the *ad libitum* and food-deprived treatments, nor between the high and low-protein artificial diets (*p > 0.05* for all comparisons; Figure 3AS and 3BS), although there was a non-significant trend for moths raised on the high-protein diet to have more IBMP than those raised on the low-protein diet. The measure of variability (variance) of SBMP (Bartlett’s K-squared= 5.9032, df = 3, *p= 0.1164*) did not differ among the different diets, whereas IBMP (Bartlett’s K-squared= 8.0726, df = 3, *p= 0.04453*) was different among treatments. The moths raised on low-protein diet had the highest variance in the amount of IBMP, and months raised in the *ad libitum* natural diet had the lowest. The volume of thoracic fluid produced did not significantly correlate with the abundance of either pyrazine (*p > 0.05* for both). There was no correlation between the amount of SBMP and IBMP and hue, saturation and brightness of the hindwings and melanin amount of fore- and hindwings (Spearman’s rank correlation, *p > 0.05* for all comparisons).

### Predator assays

When compared to the control (water), the high-protein, and the *ad libitum* natural diets led to a more effective chemical defence fluid in female wood tiger moths. The proportion of bait eaten was higher when the birds ate baits soaked in water than baits soaked in fluid from moths raised on the high-protein diet (z = -2.91, *p = 0.004*, Figure 2A) and *ad libitum* natural diet (z = -2.83, *p = 0.005*, Figure 2A). No significant differences were found from the water control for the other dietary treatments (*p>0.05* for all values, see Supplementary Material, Table 11S). Birds cleaned their beak more frequently when they were exposed to the fluid from moths raised in high-protein diet (z = 2.26, *p = 0.02*, Figure 2C) and *ad libitum* natural diet (z = 2.04, *p = 0.04*, Figure 2C) compared to water, but not when they ate the fluid from moths reared in low-protein diet and food deprivation treatments (*p>0.05* for all the comparisons, see Supplementary Material, Table 19S). Likewise, we found that the latency to approach was higher than the water control for the fluids of moths from the high-protein diet (z = -2.47, *p = 0.013*) and to a lesser extent the *ad libitum* natural diet (z = -1.91, *p = 0.056*). Similarly, the latency to eat was longer than the control when the predators approach the bait of females raised in high-protein diet (z = -2.45, *p = 0.014*). There were no differences in the latency to eat after approaching when the birds were fed with baits soaked in fluid from moths reared in all the different treatments compared to water (*p>0.05* for all the values, see Supplementary Material, Table 15S). Both natural food deprivation and protein-limited diet made defence fluids less unprofitable suggesting that moths invest more to chemical defence when resources are high. The proportion of bait eaten (a measure of palatability) was significantly lower for fluids of female moths fed with *ad libitum* natural diet compared to those that were food deprived (z=-2.29 *p= 0.02,* Figure 2A) with a similar trend for the fluids of moths raised in low-protein diet to be eaten faster than high-protein ones (z=-1.91, *p=0.056,* Figure 2A). There was no significant overall difference between the natural and artificial diets (z=-0.44, *p=0.66*, Figure 2A). The latency to approach differed between high and low-protein diet (z=-3.74, *p=<0.001*, Figure 2B), while the latency to eat after approaching and latency to eat the bait showed no significant differences between the chemical defence of moths raised in different treatments (*p>0.05* for all values, see Supplementary Material, Table 16S, 18S). Nor did the beak wiping behaviour (*p>0.05* for all values; Figure 2C, see Supplementary Material, Table 20S).

**Figure 2.**
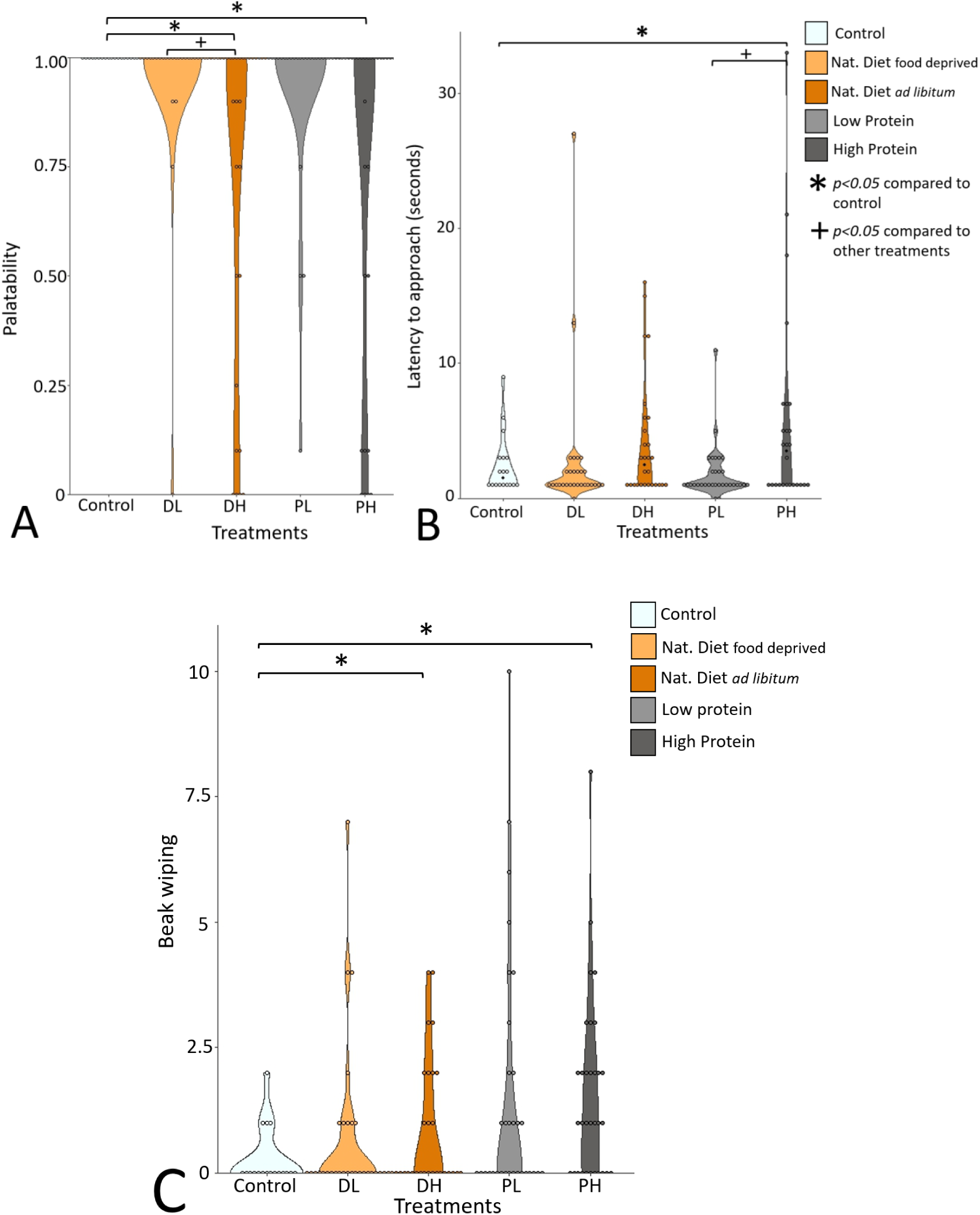
**A)** Palatability (proportion of bait soaked in female moths’ fluid eaten). **2 B)** Latency to approach (in seconds) the bait soaked in female moths’ fluid. **2 C)** Beak wiping events per minute in response to baits soaked in female moths’ fluid. (DL = natural diet food-deprived, DH = natural diet *ad libitum*, PL = Low-protein artificial diet, PH = High-protein artificial diet)

We found a correlation between the palatability (proportion of bait eaten) of the chemical defence fluid and the hue in females raised on the high-protein diet: moths with higher hue values (redder; Spearman’s rank correlation, rs = -0.5411*; p = 0.04*, Figure 3) and higher saturation (Spearman’s rank correlation, rs = -0.55; *p = 0.03*, Figure 3) hindwings, have more deterrent chemical defences. However, this was not the case for moths raised on the low-protein diet. There was no correlation between the effectiveness of the defences and the melanin amount of fore- and hindwings in moths raised in high and low-protein diet treatments (*p > 0.05* for all comparisons).We found no correlation between the efficacy of the chemical defences and hue, saturation and brightness of the hindwings and melanin amount of fore- and hindwings in moths raised in *ad libitum* and food-deprived natural diets, however the sample size was relatively small (*ad libitum* diet, *n =* 10, and food-deprived diet, *n =* 11; *p > 0.05* for all comparisons; Figure 4). We found no correlation between the palatability and the UV and red channel values across treatments (*p > 0.05* for all comparisons). There was no significant correlation between palatability and the blue and green channel values across treatments (*p > 0.05* for all comparisons), nor in the luminance (*p > 0.05* for all comparisons), except for the high-protein diet group. However, high blue channel values from individuals raised in high protein diet were significantly associated with lower palatability (Spearman’s rank correlation, rs = 0.89; *p = 0.04*). Likewise, high green channel values (Spearman’s rank correlation, rs = 0.90; *p = 0.05*) and high luminance values (Spearman’s rank correlation, rs = 0.85; *p = 0.04*) in the wings were associated with lower palatability in individuals raised in high protein diet.

**Figure 3.**
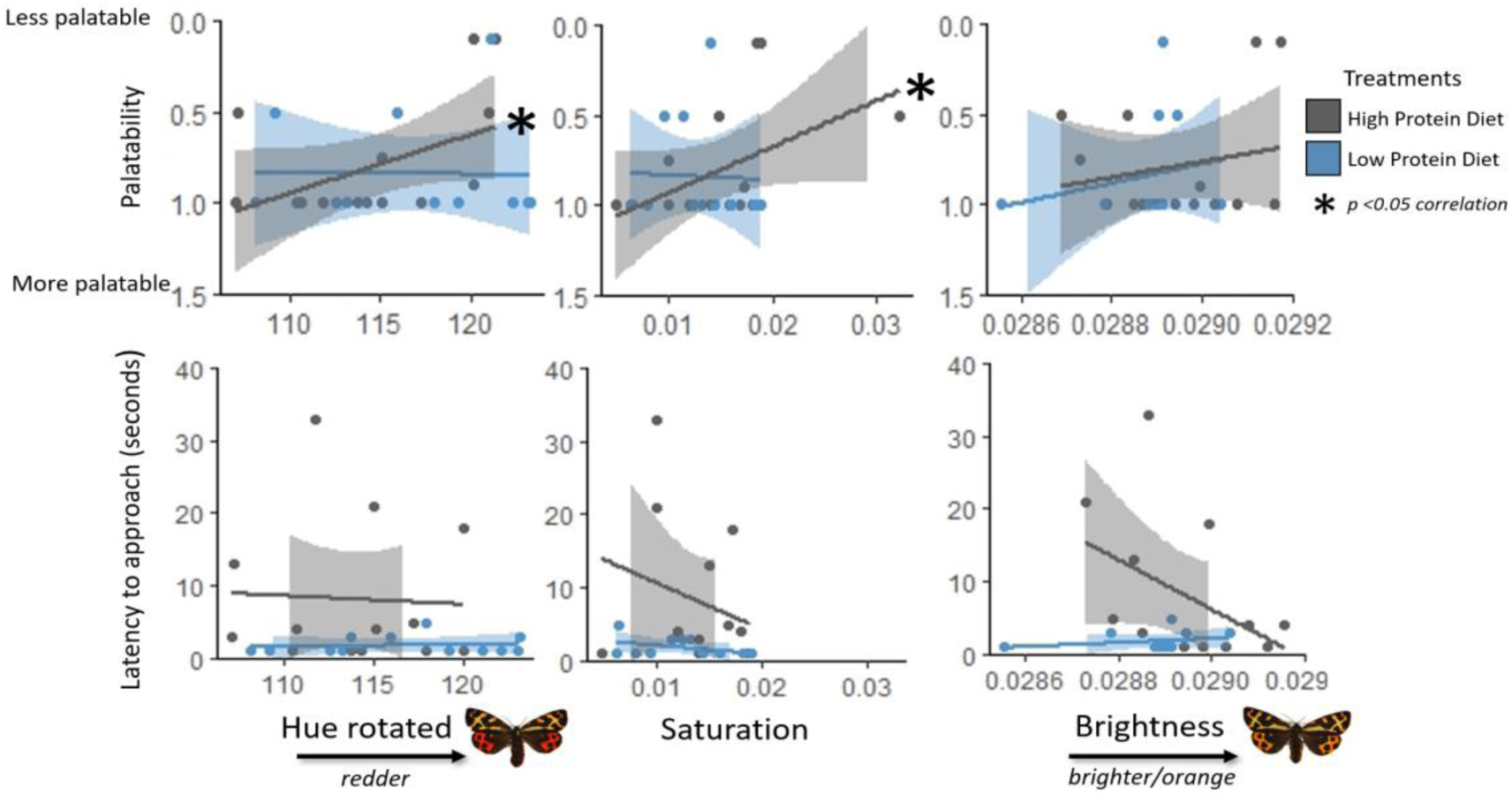
Top: correlation between the proportion of bait soaked in female moths’ fluid (raised in high-protein, *n = 15*, and low-protein, *n = 12*, diets) eaten by predators (palatability) with hue, saturation and brightness. Bottom: correlation between the latency to approach (in seconds) the bait soaked in female moths’ fluid (raised in high-protein, *n = 15*, and low-protein, *n = 12,* diets) with hue, saturation and brightness.

**Figure 4.**
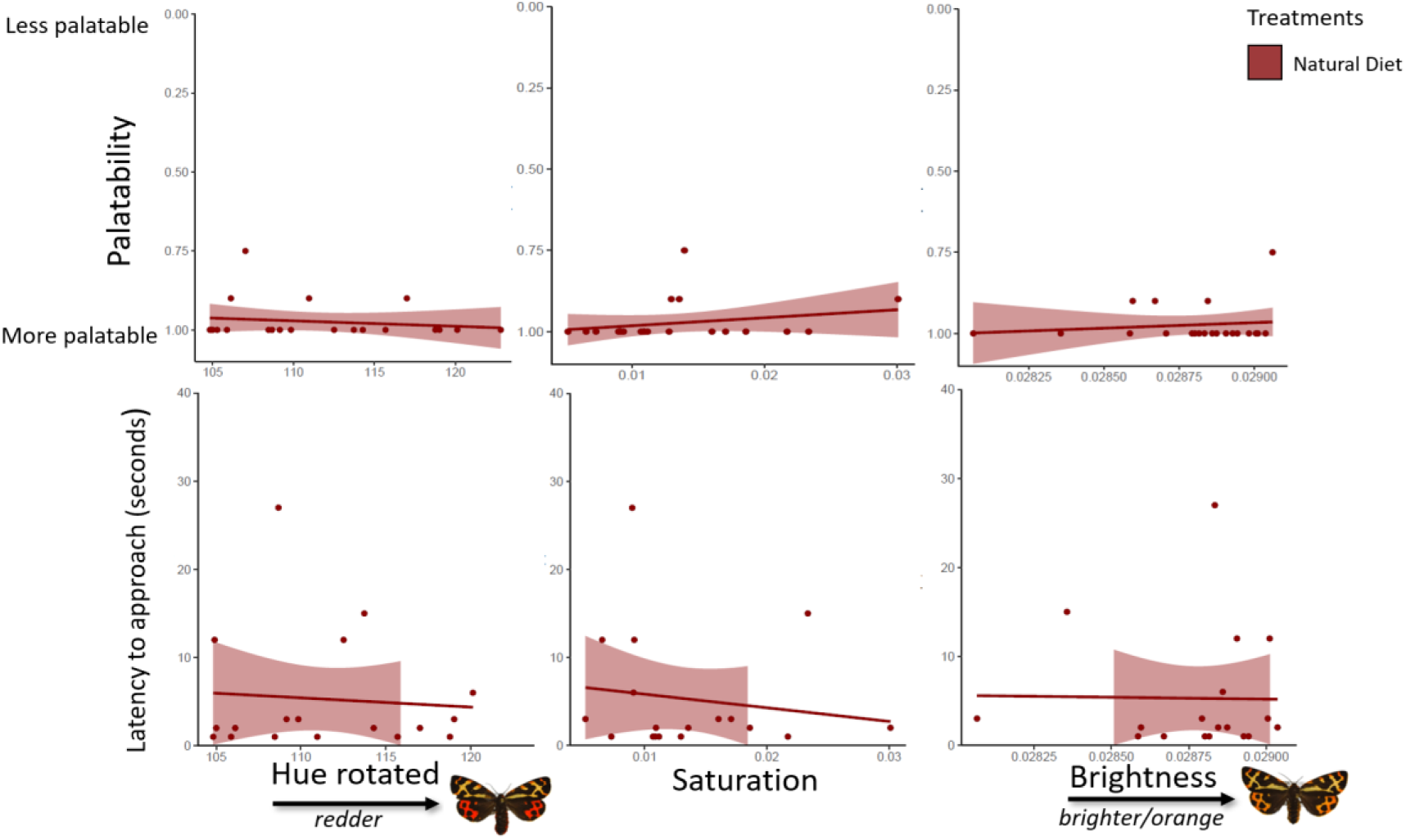
Top: correlation between the proportion of bait soaked in female moths’ fluid (raised in both *ad libitum, n =* 10, and food-deprived, *n =* 11, diets) eaten by predators (palatability) with hue, saturation and brightness. Bottom: correlation between the latency to approach (in seconds) the bait soaked in female moths’ fluid (raised in both *ad libitum, n = 10,* and food deprived, *n = 11,* diets) with hue, saturation and brightness.

## Discussion

Contrary to our expectations, we found that high resource availability during development (a high-protein diet) resulted in a positive (“honest”) correlation between primary and secondary defences in the aposematic wood tiger moth. The efficacy of the defensive fluid, as well as the saturation of the wings, increased with the proteins’ availability in the larval diet. However, on all other dietary treatments, this correlation was not found. This emphasises the importance of considering multiple levels of resource availability when trying to measure correlations between warning signals and defence, particularly in laboratory experiments, as they may not reflect the reality in the wild.

### Larval resources influence chemical defence against predators

In keeping with Burdfield-Steel et al. (2019), we found that individuals raised on a natural diet had less effective chemical defences when food-deprived once per week. While comparisons between the natural and the artificial diet treatments are difficult to interpret, given both the nutritional differences between the two types of diets (see Ojala et al., 2005 for more details) and the life-history changes they induced, the differences between the control and food-deprived treatments and the high- and low-protein artificial diets show that both short-term food deprivation and a reduction in protein availability negatively affected the insects’ defences, even when the toxins are not sequestered but produced *de novo* (Zvereva and Kozlov, 2016).

In wood tiger moths the methoxypyrazine SBMP has previously been shown to be deterrent to birds, with its effects increasing with concentration (Rojas et al., 2017, Ottocento et al., 2022). In contrast, IBMP had reduced effects on birds at higher concentrations, but in combination with SBMP at a 50/50 ratio influences a greater number of birds’ behaviour (e.g., the proportion of bait eaten decreased, and increase the drinking behaviour) than SBMP alone, highlighting the synergistic, rather than additive, effect of the combination of these two pyrazines (Ottocento et al., 2022). We predicted that moths raised on high-protein and *ad libitum* natural diet will have higher levels of SBMP and IBMP. In this study, we found that moths raised on high- and low-protein artificial diets had higher levels of both pyrazines (SBMP and IBMP) in the defensive fluid than the natural diets, perhaps because these diets were created to aid in the rearing of Lepidoptera in the laboratory and may contain higher amounts of key micronutrients (Morehouse & Rutowski, 2010). While we did not detect significant differences in the abundance of the SBMP and IBMP between the high- and low-protein diets (and between food-deprived and *ad libitum* natural diet treatments), we did see a trend of higher amounts of methoxypyrazines, in particular IBMP in moths from the high-protein treatment. We hypothesised that moths raised in high-protein and *ad libitum* natural diets would have the lower variance in pyrazines SBMP and IBMP. While we did not find any difference in variance among treatments in SBMP, we found that the highest variances in IBMP abundance were in moths from the low-protein treatment. It is possible that in a scenario of lack of nutrients, the production of IBMP is not prioritised, while the production of SBMP (which is more deterrent) is maintained even under resource stress.

More importantly, the bird response data showed that both the reduced protein diet and the natural deprivation diet did result in detectable reductions in the effectiveness of the defensive fluids to repel an ecologically relevant predator (blue tits). These results agree with previous studies that found that the Estonian male wood tiger moths raised on high-protein diets have the most deterrent chemical fluid (Ottocento et al., 2023) and that food-deprived moths (fed on natural diet) have fewer repellent defences (Burdfield-Steel et al., 2019). Not surprisingly, nitrogen, the core component of amino-acids and thus proteins, is essential in the biosynthesis of pyrazines (Hodge, Mills & Fisher, 1972; Wong & Bernhard, 1988). Therefore, variation in the protein available in the diet directly affects pyrazine production and hence chemical defence (Ottocento et al., 2023).

### Protein availability, but not food deprivation affects warning signals

When looking at the aposematic hindwing colouration we hypothesised that moths from the high-protein and *ad libitum* treatments would have the most conspicuous colouration (such as redder hue, higher saturation, lower brightness, and higher red colour channel values) compared to those raised on the low-protein diet. Indeed, we found that female moths raised on a low-protein diet displayed a decrease in saturation and an increase in the brightness of the hindwings when compared to those raised on a high-protein diet, thus being paler and duller. Saturation (signal intensity) has been shown in many systems to play an important role in the effectiveness of warning colouration, and the reduction in saturation and the increase in brightness (which results in pale orange hindwings) in moths from low-protein diet, points to a weaker visual signal. High resource conditions did not result in a significant increase in hue (redness). Previous research has demonstrated that darker, redder hindwings are a stronger warning signal, and this difference in conspicuousness is detectable by avian predators (Lindstedt et al., 2011). Using the avian vision model, we found that moths raised on both artificial protein diets showed a slight, although non-significant, increase in the red and green channel values compared to those on natural diets (suggesting that the artificial supplements can impact production of the pigments responsible for the visual warning signal), but this difference was not noticeable between high-low protein diets and *ad libitum* and food deprived treatments. This is particularly interesting as previous studies found high heritability in the production of red pigments (pheomelanin; M. Brien, personal communication) in *A. plantaginis* (Lindstedt et al., 2016), thus, the genetic variation may overshadow any environmentally induced variation in this trait. These results support previous research showing that food deprivation, in the context of a natural diet, did not affect female hindwing colouration (Burdfield-Steel et al., 2019). The colour properties that were significant in the HSV colour space, however, were not confirmed by the avian vision model channel values, which warrants further investigation.

We predicted that individuals raised on the *ad libitum* natural diet and high-protein diet would have greater amounts of melanin than individuals raised on the food-deprived natural diet and low-protein diet. The amount of melanin in the fore- and hindwings did not differ within treatments, but did vary between artificial and natural diets: individuals raised on artificial diets had more melanin pigmentation than the moths reared on natural diets. Melanin is strongly affected by the protein quality of the diet in insects (Lee et al., 2008) and while we did not find a difference between high- and low-protein treatments, it is possible that the artificial diet contains macronutrients that are not present in the natural diets.

The phenomenon of automimicry may also explain why the warning signal (composed of both the red and black colouration in the wings) may be less affected by dietary changes than other traits such as chemical defences: within a population, it is possible that some individuals lack chemical defences but are protected from predators by their resemblance to more strongly defended members of the same species (Brower et al., 1967; Speed et al., 2012). Indeed, the availability of resources in the wild can vary and lead to automimicry (Ruxton, Sherratt & Speed, 2019; Speed et al., 2012), and this study can be viewed as an additional example of that.

### Larval resources influence life-history traits

By manipulating the diet of female wood tiger moths, we demonstrate that resource availability significantly influenced larval life-history. In keeping with our hypothesis, there was a trend of increasing time to develop from natural to artificial diets, and from high to low protein diet and *ad libitum* to food-deprived treatments. A similar pattern was found in male moths (Ottocento et al., 2023), suggesting food deprivation and lack of nutrients slow down the development process. In addition, larvae fed on the natural diet often hid in rolled-up leaves to protect themselves during the pupation process, while those fed on the artificial diet did not have such hiding spots. This difference may have led to a reduced stimulus for eating. We did not however see significant differences in pupal weight or wing size between the *ad libitum* and food-deprived diets, or between the high and low protein diets. In fact, pupal weight and the forewing size were greater in moths raised on artificial diet compared to those raised on natural diet, which may indicate that despite the slower developmental time, the larvae had more time to acquire resources and grow.

### High resource availability, but not low, leads to honest signals

We found a positive correlation between the strength of signal and defence only in the moths raised on the high protein diet. While comparison between artificial and natural diets are difficult given that the former may contain micronutrients missing from the latter, the high protein diet clearly represents a higher resource diet for the wood tiger moth than the low protein, as females raised on it not only had stronger warning signals and chemical defences but also laid more eggs.

Interestingly, we found correlations between the unpalatability of the chemical defences and the blue and green colour channels in individuals raised on a high-protein diet, suggesting that an increase in resource availability affects a part of the signal not typically associated with aposematic effectiveness. In many aposematic organisms, reliable signals of toxicity are commonly linked to red pigments which are detectable by avian predators and often associated with unpalatability (Thery and Gomez 2010; Stevens and Ruxton 2012). The unexpected association with the blue and green colour channels may reflect a resource-driven enhancement of pigmentation that does not necessarily indicate toxicity but might still increase the signal conspicuousness, thereby affecting predator response.

These results support one of the underlying assumptions of the resource allocation hypothesis, as manipulation of a single resource, in this case, protein in the diet, caused changes in both the strength of warning colouration and chemical defence - and influenced predators’ behaviour. However, they cast doubt on whether the relationship between hindwing colouration and chemical defence in wild wood tiger moths is likely to be consistent enough to be of use to predators. Indeed, we did not find any correlation between the effect of the chemical defences on predators and the warning signal in moths raised in the two natural diets (*ad libitum* and food deprivation). Taken together the changes in the colour seen in this study suggest that variation in resources can create a positive relationship between the effectiveness of the chemical defence and saturation in the hindwings, but only under certain circumstances.

It is also important to consider that the cost of conspicuousness may vary when the individuals sequestered their toxins rather than producing them *de novo*. The sequestration process (and subsequent detoxification) may require an extra expenditure of energy in processing and eliminating toxins which might lead to a weakened warning signal (Lindstedt et al., 2010b). Such alterations in diet quality are important in polyphagous species and both food deprivation and detoxification may yield distinct impacts on the wing’s phenotype (Burdfield-Steel et al., 2019). Lindstedt et al., (2010) found that female wood tiger moths raised in high concentrations of iridoid glycosides present weaker warning signals due to the high excretion costs of the toxins ingested. A recent study focusing on the monarch butterfly *Danaus plexippus*, revealed a connection between toxin sequestration and warning signals: males with more prominent warning signals had lower levels of oxidative damage, which is caused by higher concentrations of sequestered cardenolides (Blount et al., 2023). In the large milkweed bug, *Oncopeltus fasciatus* a greater sequestration of cardenolides led to a lower amount of the antioxidant glutathione, which translates to lower luminant orange warning signals compared to the animals that have more glutathione (Heyworth et al., 2023). Indeed, when wood tiger moths were allowed to sequester pyrrolizidine alkaloids (which were not present in any of the dietary treatments in this study), this was found to negatively trade-off with defence against the early stages of predator attack, suggesting that it reduces investment in pyrazines (Winters et al., 2021). Thus, in species able to both sequester and produce chemical defences *de novo* (e.g., the red postman *Heliconius erato* see Mattila et al., 2021; the burnet moth *Zygaena filipendulae* see Fürstenberg-Hägg et al., 2014; the wood tiger moths *A. plantaginis*, see Burdfield-Steel et al., 2018) the mechanisms underlying investment into the different components of aposematism may be even more complex.

## Conclusion

In conclusion, this study found evidence that resource availability, and subsequent allocation, can impact both warning signals and chemical defence in the wood tiger moth - as well as the relationship between the two. This finding highlights the importance of establishing if observed variation in chemical defences are detectable, and thus subject to selection by, ecologically relevant predators. In addition, it emphasises the need to consider different levels of resource availability in studies of honest signalling. If the pattern we found is shared across species, then studies of laboratory populations, where resources are often abundant and consistent, may find evidence of honest signalling, even if it does not occur in wild populations. Thus, studies looking for honest signalling in laboratory populations may be in danger of overestimating its prevalence.

## Author contributions

EBS, JM and BR conceived the study. MF carried out the rearing experiments, and MF, ON and CO performed the colour analysis. EBS performed the chemical analysis. CO and BR did the bird experiments. EBS, MF and CO did the data analysis. EBS and CO wrote the first draft, and all authors contributed to the final manuscript.

## Acknowledgments

We are grateful to Kaisa Suisto and the greenhouse workers at the University of Jyväskylä for the rearing of the laboratory moths and Helinä Nisu assisted with the catching and training of wild birds. We thank Konnevesi Research Station for the facilities. We also thank David Shuker for comments on an earlier version of this manuscript.

## Funding

This study was funded by the Academy of Finland the project No. 345091 (to JM).

## Notes

### Competing Interest Statement

The authors have declared no competing interest.

### Summary of Updates

Update the abstract of the manuscript

