## Supplementary Material for "Honest signalling in predator-prey interactions: testing the resource allocation hypothesis"

#### 1. Protein diet

**Table 1S** High and low protein diet

| **High protein artificial diet** | **Low protein artificial diet** |
| --- | --- |
| Agar g 18 | Agar g 18 |
| Wheat Germ g 72 | Wheat Germ g 72 |
| **Teklad VitFree Casein g 32.4** | **Teklad VitFree Casein g 10.8** |
| Sucrose g 28.8 | Sucrose g 28.8 |
| Salt Mix g 10.8 | Salt Mix g 10.8 |
| Torula Yeast g 14.4 | Torula Yeast g 14.4 |
| **Cellulose g 0** | **Cellulose g 21.6** |
| Cholesterol g 3.15 | Cholesterol g 3.15 |
| Sorbic Acid g 1.8 | Sorbic Acid g 1.8 |
| Ascorbic Acid g 3.6 | Ascorbic Acid g 3.6 |

| | |
|---|---|
| VanderZandt Vit Mix g 12.6 | VanderZandt Vit Mix g 12.6 |
| Water (cool) ml 450 | Water (cool) ml 450 |
| Water (boiling) ml 450 | Water (boiling) ml 450 |
| Linseed oil ml 6 | Linseed oil ml 6 |

**Important note:**

The casein used is Teklad Vitamin-Free. In this diet, casein is the predominant source of protein. If another type of casein is used, the amount of vitamins may also vary.

#### 2. Life history

##### 2.1 Developmental time per day; pupae weight

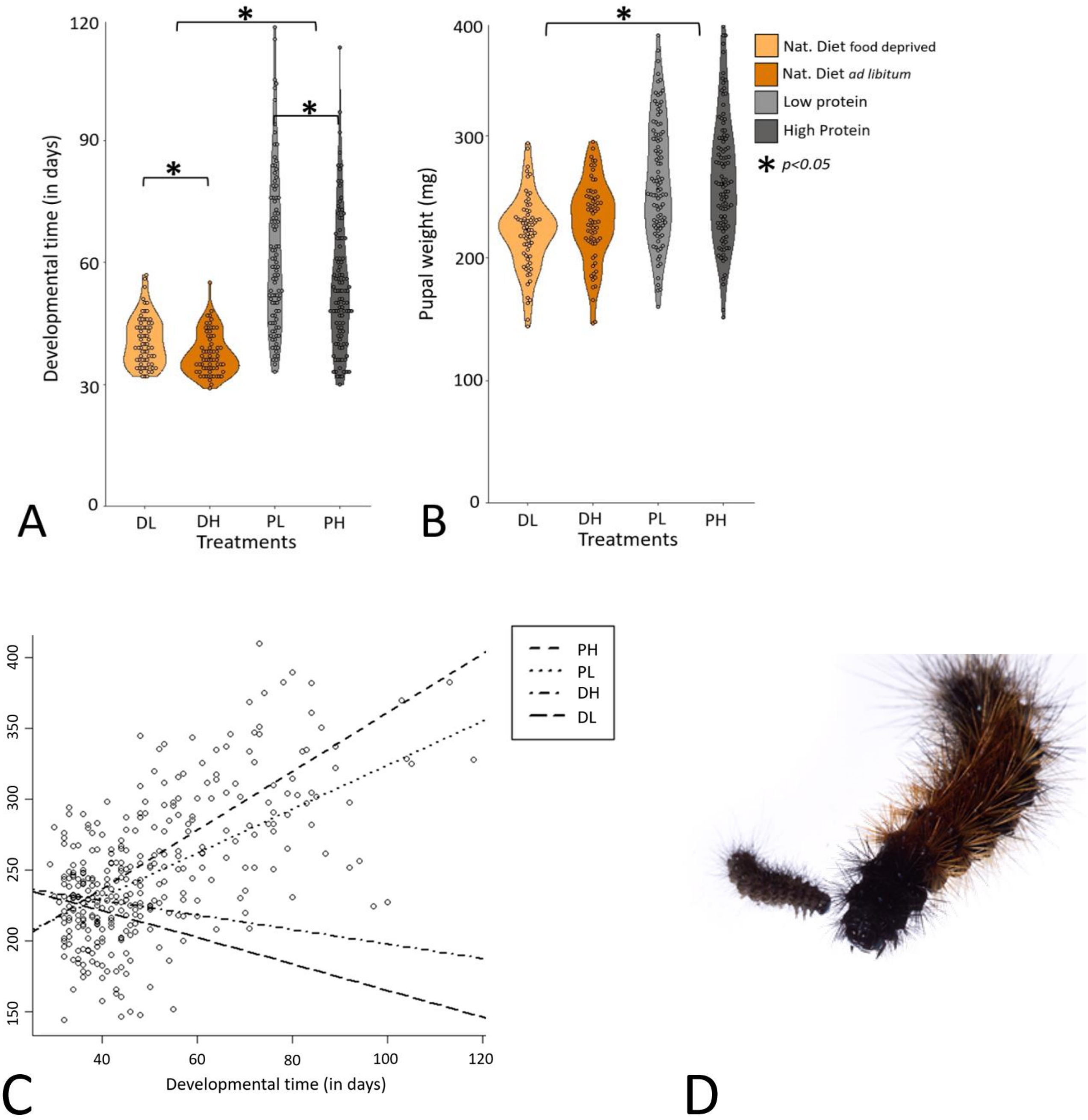


**Supplementary Figure 1**. Top row: the effect of the four dietary treatments (DL = natural diet food-deprived, DH = natural diet *ad libitum*, PL = Low-protein artificial diet, PH = High-protein artificial diet) and development time in days (A) and pupa weight in milligrams (B) of female moths. Asterisks indicate significant differences ($p < 0.05$). Bottom row: the correlation between development time and pupa weight (C). Lines indicate each of the four dietary treatments. Photo of wood tiger moth larva (D) courtesy of Sebastiano De Bona.

**Table 2S** The difference in developmental time between *ad libitum* and food-deprived natural diets and low- and high-protein diets.

| | coef | exp (coef) | se (coef) | z | p |
|---|---|---|---|---|---|
| ***Ad libitum*-Food depriv.** | -0.33 | 0.72 | 0.09 | -3.65 | 0.0003 |
| **Natural Diet-Artificial Diet** | -1.11 | 0.33 | 0.08 | -13.48 | 0.0000 |
| **High Protein-Low Protein** | -0.25 | 0.77 | 0.075 | -3.40 | 0.0007 |

**Table 3S** The difference in pupae weight between *ad libitum* and food-deprived natural diets and low- and high-protein diet.

| | Value | Std. Error | t value | p value |
|---|---|---|---|---|
| **Intercept** | 5.44 | 0.03 | 168.024 | <2e-16 *** |
| ***Ad libitum*-Food depriv.** | -0.03 | 0.01 | -2.08 | 0.0377 * |
| **Natural Diet-Artificial Diet** | -0.088 | 0.009 | 9.39 | <2e-16 *** |
| **High Protein-Low Protein** | 0.004 | 0.011 | 0.35 | 0.73 |

#### 3.1 Female fecundity

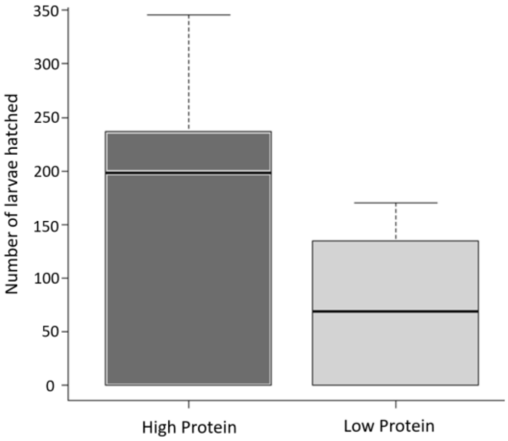


**Supplementary Figure 2.** The number of hatched larvae produced by females raised on high- and low-protein diets.

### 4. Colour measurements

#### 4.1 Hue

**Table 4S** The difference in hue between diet treatments

| | Value | Std. Error | DF | t value | p value |
|---|---|---|---|---|---|
| **Intercept** | 118.43 | 1.28 | 87 | 92.74 | 0.00 |
| ***Ad libitum*-Food depriv.** | -0.18 | 0.64 | 87 | -0.28 | 0.78 |
| **Natural Diet-Artificial Diet** | -0.60 | 0.47 | 87 | 1.27 | 0.21 |
| **High Protein-Low Protein** | -0.62 | 0.57 | 87 | -1.08 | 0.28 |

#### 4.2 Saturation

**Table 5S** The difference in saturation between diet treatments

| | Value | Std. Error | DF | t value | p value |
|---|---|---|---|---|---|
| **Intercept** | 0.62 | 0.007 | 87 | 83.74 | 0.00 |
| ***Ad libitum*-Food depriv.** | 0.007 | 0.007 | 87 | 1.06 | 0.29 |
| **Natural Diet-Artificial Diet** | 0.001 | 0.005 | 87 | 0.28 | 0.78 |
| **High Protein-Low Protein** | 0.02 | 0.007 | 87 | 3.10 | 0.003 |

#### 4.3 Brightness

**Table 6S** The difference in brightness between diet treatments

| | Value | Std. Error | DF | t value | p value |
|---|---|---|---|---|---|
| **Intercept** | 0.41 | 0.004 | 87 | 96.74 | 0.00 |
| ***Ad libitum*-Food depriv.** | 0.004 | 0.004 | 87 | 1.08 | 0.28 |
| **Natural Diet-Artificial Diet** | 0.016 | 0.003 | 87 | 5.55 | 0.00 |
| **High Protein-Low Protein** | -0.01 | 0.004 | 87 | -3.00 | 0.004 |

#### 4.4 Hindwing Melanin

**Table 7S** The difference in the hindwing melanin between diet treatments

| | Value | Std. Error | DF | t value | p value |
|---|---|---|---|---|---|
| **Intercept** | 0.64 | 0.01 | 87 | 52.52 | 0.00 |
| ***Ad libitum*-Food depriv.** | -0.01 | 0.009 | 87 | -1.52 | 0.13 |
| **Natural Diet-Artificial Diet** | 0.01 | 0.006 | 87 | 3.00 | 0.0035 |
| **High Protein-Low Protein** | -0.002 | 0.008 | 87 | -0.24 | 0.81 |

#### 4.5 Forewing Melanin

**Table 8S** The difference in the forewing melanin between diet treatments

| | Value | Std. Error | DF | t value | p value |
|---|---|---|---|---|---|
| **Intercept** | 0.58 | 0.008 | 87 | 73.04 | 0.00 |
| ***Ad libitum*-Food depriv.** | -0.008 | 0.006 | 87 | -1.29 | 0.20 |
| **Natural Diet-Artificial Diet** | 0.01 | 0.004 | 87 | 2.66 | 0.0094 |
| **High Protein-Low Protein** | 0.003 | 0.005 | 87 | 0.59 | 0.55 |

#### 4.6 Hindwing area

**Table 9S** The difference in the hindwing area between diet treatments

| | Value | Std. Error | DF | t value | p value |
|---|---|---|---|---|---|
| **Intercept** | 88528.92 | 2259.18 | 87 | 39.19 | 0.00 |
| ***Ad libitum*-Food depriv.** | 458.74 | 1486.24 | 87 | 0.31 | 0.76 |
| **Natural Diet-Artificial Diet** | 1624.83 | 1080.19 | 87 | 1.50 | 0.14 |
| **High Protein-Low Protein** | 1384.32 | 1331.87 | 87 | 1.04 | 0.30 |

#### 4.7 Forewings area

**Table 10S** The difference in the forewing area between diet treatments

| | Value | Std. Error | DF | t value | p value |
|---|---|---|---|---|---|
| **Intercept** | 105901.69 | 2276.10 | 87 | 46.53 | 0.00 |
| ***Ad libitum*-Food depriv.** | 6.45 | 1482.50 | 87 | 0.004 | 0.99 |
| **Natural Diet-Artificial Diet** | 2373.12 | 1078.12 | 87 | 2.20 | 0.03 |
| **High Protein-Low Protein** | 1286.86 | 1328.51 | 87 | 0.97 | 0.33 |

#### 4.8 U channel

**Table 11S** The difference in the ultraviolet values between diet treatments

| | Estimate | Std. Error | DF | t value | p value |
|---|---|---|---|---|---|
| **Intercept** | 17898,94 | 757,053 | 43 | 23,643 | 6.45e-05 |
| ***Ad libitum*-Food depriv.** | -246,517 | 1039,04 | 43 | -0,237 | 0,814 |
| **Natural Diet-Artificial Diet** | 465,882 | 689,428 | 43 | 0,676 | 0,503 |
| **High Protein-Low Protein** | 155,269 | 911,368 | 43 | 0,17 | 0,866 |

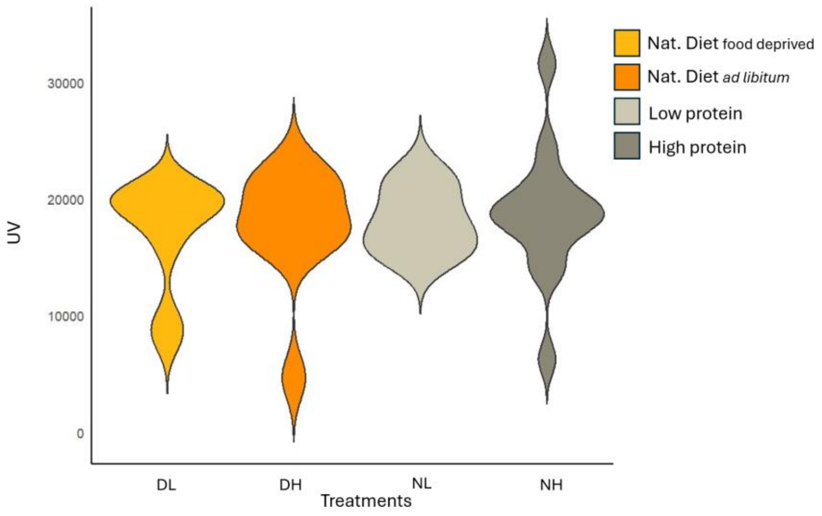


**Supplementary Figure 3.** UV measure of female moths' wings from the four different diets. (DL = natural diet food-deprived, DH = natural diet *ad libitum*, PL = Low-protein artificial diet, PH = High-protein artificial diet)

#### 4.9 B channel

**Table 12S** The difference in the blue channel values between diet treatments

| | Estimate | Std. Error | DF | t value | p value |
|---|---|---|---|---|---|

| | | | | | |
|---|---|---|---|---|---|
| **Intercept** | 18445,3 | 629 | 43 | 29,323 | <2e-16 |
| ***Ad libitum*-Food depriv.** | -476,6 | 951 | 43 | -0,501 | 0,619 |
| **Natural Diet-Artificial Diet** | 424,4 | 629 | 43 | 0,675 | 0,504 |
| **High Protein-Low Protein** | -450 | 823,6 | 43 | -0,546 | 0,588 |

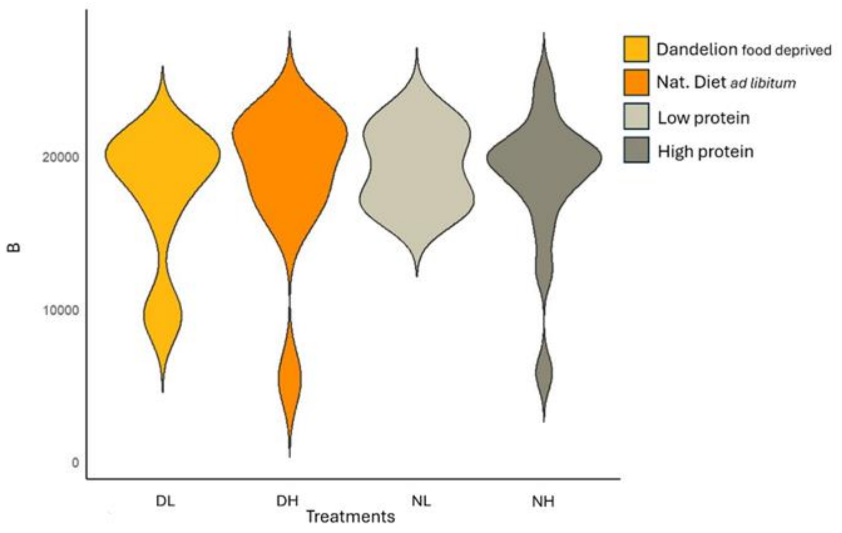


**Supplementary Figure 4.** Blue channel colour measure of female moths' wings from the four different diets. (DL = natural diet food-deprived, DH = natural diet *ad libitum*, PL = Low-protein artificial diet, PH = High-protein artificial diet)

#### 4.10 G channel

**Table 13S** The difference in the green channel values between diet treatments

| | Estimate | Std. Error | DF | t value | p value |
|---|---|---|---|---|---|

| | | | | | |
|---|---|---|---|---|---|
| **Intercept** | 21383,08 | 500,89 | 43 | 42,691 | <2e-16 |
| ***Ad libitum*-Food depriv.** | -503,52 | 757,27 | 43 | -0,665 | 0,5097 |
| **Natural Diet-Artificial Diet** | 1004,89 | 500,89 | 43 | 2,006 | 0,0511 |
| **High Protein-Low Protein** | -15,16 | 655,81 | 43 | -0,023 | 0,9817 |

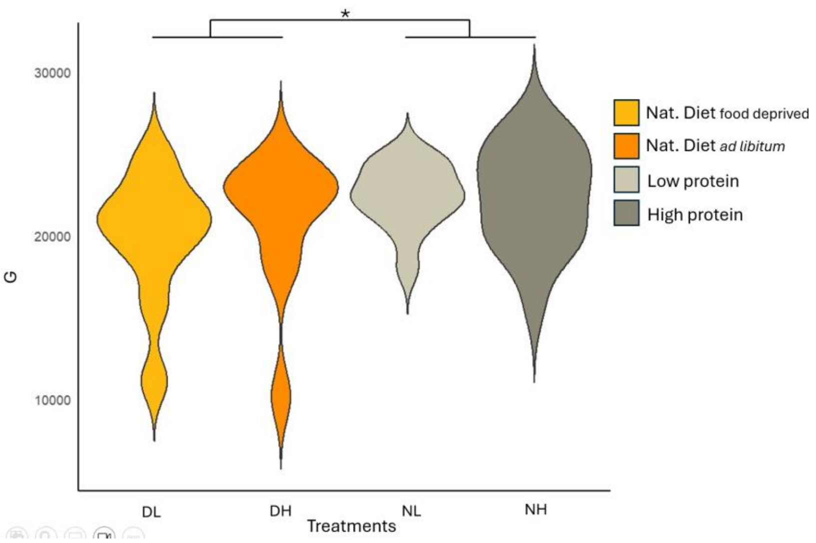


**Supplementary Figure 5.** Green channel colour measure of female moths' wings from the four different diets. (DL = natural diet food-deprived, DH = natural diet *ad libitum*, PL = Low-protein artificial diet, PH = High-protein artificial diet)

#### 4.11 R channel

**Table 14S** The difference in the red channel values between diet treatments

| | Estimate | Std. Error | DF | t value | p value |
|---|---|---|---|---|---|

| | | | | | |
|---|---|---|---|---|---|
| **Intercept** | 24529 | 502,1 | 43 | 48,848 | <2e-16 |
| ***Ad libitum*-Food depriv.** | -740,9 | 759,2 | 43 | -0,976 | 0,3346 |
| **Natural Diet-Artificial Diet** | 949,8 | 502,1 | 43 | 1,891 | 0,0653 |
| **High Protein-Low Protein** | 203,9 | 657,5 | 43 | 0,31 | 0,758 |

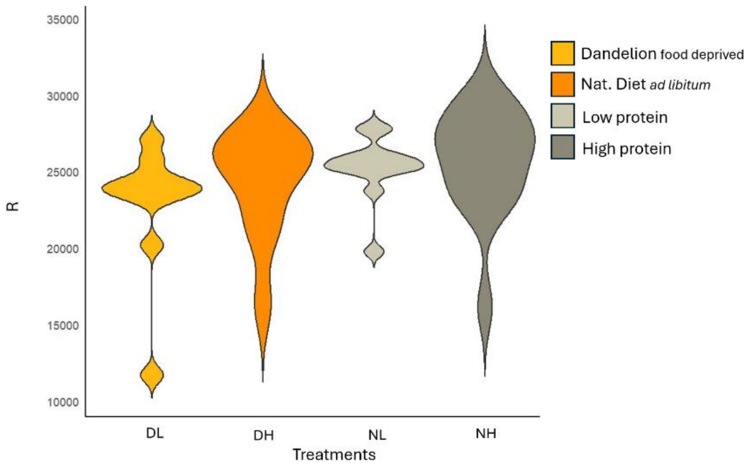


**Supplementary Figure 6.** Red channel measure of female moths' wings from the four different diets. (DL = natural diet food-deprived, DH = natural diet *ad libitum*, PL = Low-protein artificial diet, PH = High-protein artificial diet)

#### 4.12 Luminance

**Table 15S** The difference in the luminance values between diet treatments

| | Estimate | Std. Error | DF | t value | p value |
|---|---|---|---|---|---|
| **Intercept** | 22013,53 | 496,74 | 43 | 44,316 | <2e-16 |
| ***Ad libitum*-Food depriv.** | -597,19 | 751 | 43 | -0,795 | 0,4309 |

| | | | | | |
|---|---|---|---|---|---|
| **Natural Diet-Artificial Diet** | 918,83 | 496,74 | 43 | 1,85 | 0,0712 |
| **High Protein-Low Protein** | 1,24 | 650,38 | 43 | 0,002 | 0,9985 |

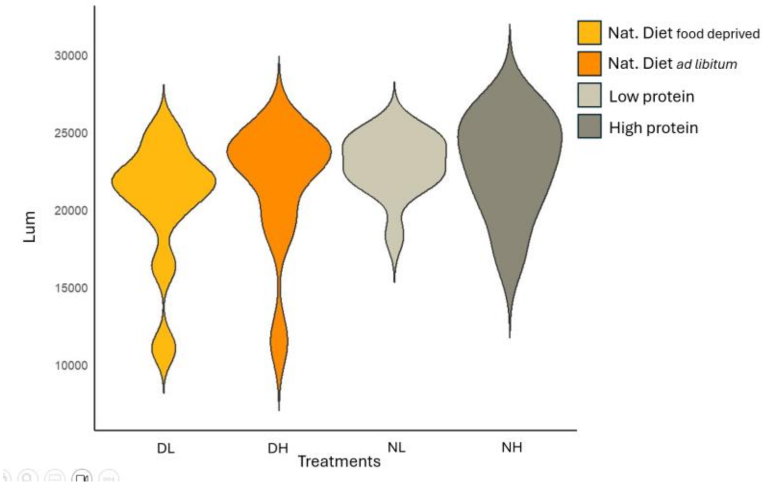


**Supplementary Figure 7.** Luminance measure of female moths' wings from the four different diets. (DL = natural diet food-deprived, DH = natural diet *ad libitum*, PL = Low-protein artificial diet, PH = High-protein artificial diet)

#### 5. Quantification of chemical defences

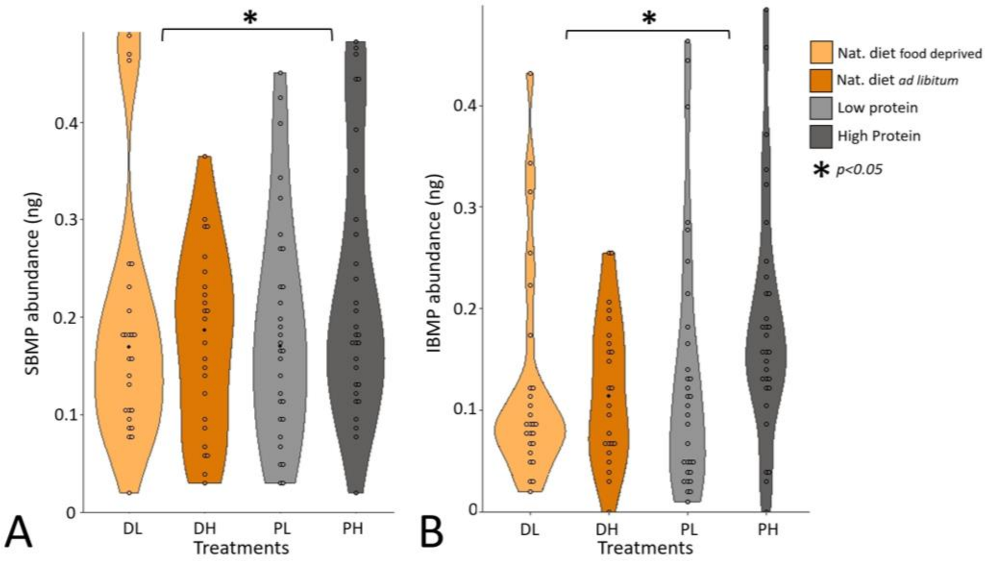

**Supplementary Figure 3.** The abundance (in ng) of the SBMP methoxypyrazine (A) and IBMP methoxypyrazine in the thoracic fluids of female moths reared on the different diets.

#### 6. Predator response to female's defensive fluid

##### 6.1 Proportion of bait eaten

**Table 11S** Proportion of bait (oat) eaten with control (water). The proportion of bait was tested using package glmmTMB with family = beta_family(link= "logit) for beta regression models with the diet treatment, and trial set as a fixed effect, bird ID as a random effect, and an offset included to account for differences in observation time.

| | Estimate | Std. Error | z value | Pr(>\|z\|) |
|---|---|---|---|---|
| **Intercept** | -0.097 | 0.77 | 0.13 | 0.90 |
| ***Ad libitum*** | -2.55 | 0.90 | -2.83 | 0.005 ** |
| **Food-deprived** | -0.69 | 0.89 | -0.77 | 0.44 |
| **High Protein** | -2.61 | 0.90 | -2.91 | 0.004 ** |
| **Low Protein** | -1.07 | 0.89 | -1.20 | 0.23 |
| **trial 3** | 0.55 | 0.19 | 2.81 | 0.005 ** |

**Table 12S** Proportion of bait (oat) eaten without control. The proportion of bait was tested using package glmmTMB with family = beta_family(link= "logit) for beta regression models with the diet treatment, the amount of melanin, and trial set as fixed effect, bird ID as random effect, and an offset included to account for differences in observation time.

| | Estimate | Std. Error | z value | Pr(>\|z\|) |
|---|---|---|---|---|
| **Intercept** | -1.98 | 0.37 | -5.38 | 7.47e-08 *** |
| **High Protein-Low Protein** | -0.74 | 0.39 | -1.91 | 0.056 |
| ***Ad libitum*-Food depriv** | -0.90 | 0.39 | 2.29 | 0.02 * |
| **Natural Diet-Artificial Diet** | -0.12 | 0.28 | -0.44 | 0.66 |
| **trial 3** | 0.65 | 0.21 | 3.07 | 0.002 ** |

#### 6.2 Latency to approach

**Table 13S** Latency to approach the bait (oat) with control (water) was tested using cox mixed-effects model using package coxme, fit by maximum likelihood. The latency to approach was set as response variable; the different types of diet (low protein, high protein) were set as predictor variables. The diet and trial were set as fixed effects, while bird ID was set as a random effect.

| | coeff | exp(coef) | se(coef) | z | p |
|---|---|---|---|---|---|
| ***Ad libitum*** | -0.65 | 0.52 | 0.34 | -1.91 | 0.056 |
| **Food-deprived** | -0.14 | 0.86 | 0.34 | -0.43 | 0.67 |
| **High Protein** | -0.87 | 0.42 | 0.35 | -2.47 | 0.013* |
| **Low Protein** | 0.32 | 1.37 | 0.33 | 0.95 | 0.34 |

**Table 14S.** Latency to approach the bait (oat) without control was tested using cox mixed-effects model using package coxme, fit by maximum likelihood. The latency to approach was set as response variable; the different types of diet were set as predictor variables. The interaction between diet (low protein, high protein), the amount of melanin and trial were set as fixed effects, while bird ID was set as a random effect.

| | coeff | exp(coef) | se(coef) | z | p |
|---|---|---|---|---|---|
| **High Protein-Low Protein** | -0.58 | 0.56 | 0.15 | -3.74 | 0.0002** |
| ***Ad libitum*-Food depriv** | 0.25 | 1.28 | 0.15 | 1.68 | 0.093 |
| **Natural Diet-Artificial Diet** | 0.06 | 1.06 | 0.10 | 0.57 | 0.57 |

#### 6.3 Latency to eat after approaching

**Table 15S** Latency to eat after approaching the bait (oat) with control (water) was tested using cox mixed-effects model using package coxme, fit by maximum likelihood. The latency to eat after approaching was set as response variable; the different types of diet (low protein, high protein) were set as predictor variables. The diet and trial were set as fixed effects, while bird ID was set as a random effect.

| | coeff | exp(coef) | se(coef) | z | p |
|---|---|---|---|---|---|
| ***Ad libitum*** | -0.17 | 0.85 | 0.43 | -0.38 | 0.70 |
| **Food-deprived** | -0.16 | 0.85 | 0.43 | -0.39 | 0.70 |

| | | | | | |
|---|---|---|---|---|---|
| **High Protein** | -0.23 | 0.79 | 0.44 | -0.54 | 0.59 |
| **Low Protein** | 0.50 | 0.61 | 0.43 | 1.17 | 0.24 |

**Table 16S.** Latency to eat after approaching the bait (oat) without control was tested using cox mixed-effects model using package coxme, fit by maximum likelihood. The latency to eat after approaching was set as response variable; the different types of diet were set as predictor variables. The interaction between diet (low protein, high protein), the amount of melanin and trial were set as fixed effects, while bird ID was set as a random effect.

| | **coeff** | **exp(coef)** | **se(coef)** | **z** | **p** |
|---|---|---|---|---|---|
| **High Protein-Low Protein** | 1.23 | 1.13 | 0.19 | 0.65 | 0.52 |
| ***Ad libitum*-Food depriv** | 0.001 | 1.00 | 0.19 | 0.01 | 1.00 |
| **Natural Diet-Artificial Diet** | -0.09 | 0.90 | 0.13 | -0.74 | 0.46 |

#### 6.4 Latency to eat

**Table. 17S** Latency to eat the bait (oat) with control (water) was tested using cox mixed-effects model using package coxme, fit by maximum likelihood.The latency to eat was set as response variable; the different types of diet (low protein, high protein) were set as predictor variables. The diet and trial were set as fixed effects, while bird ID was set as a random effect.

| | coeff | exp(coef) | se(coef) | z | p |
|---|---|---|---|---|---|
| ***Ad libitum*** | -0.79 | 0.45 | 0.49 | -1.60 | 0.11 |
| **Food-deprived** | -0.50 | 0.61 | 0.49 | -1.02 | 0.31 |
| **High Protein** | -1.23 | 0.29 | 0.50 | -2.45 | 0.014* |
| **Low Protein** | 0.40 | 0.67 | 0.49 | -0.82 | 0.41 |

**Table 18S.** Latency to eat the bait (oat) without control was tested using cox mixed-effects model using package coxme, fit by maximum likelihood. The latency to eat was set as response variable; the different types of diet were set as predictor variables. The interaction between diet (low protein, high protein), the amount of melanin and trial were set as fixed effects, while bird ID was set as a random effect.

| | coeff | exp(coef) | se(coef) | z | p |
|---|---|---|---|---|---|
| **High Protein-Low Protein** | -0.41 | 0.66 | 0.22 | -1.82 | 0.07 |
| ***Ad libitum*-Food depriv** | 0.14 | 1.15 | 0.22 | 0.63 | 0.53 |
| **Natural Diet-Artificial Diet** | -0.08 | 0.92 | 0.16 | -0.55 | 0.58 |

##### 6.5 Beak wiping

**Table 19S.** Beak wiping reaction to the bait (oat) with control (water) was tested using a generalised linear mixed-effects models (GLMM) with a log link and Poisson distribution, fit by maximum likelihood (Laplace approximation) to test for differences in beak wiping events per minute.The diet (low protein, high protein) and trial were set as fixed effects, while the beak wiping frequency was set as response variable and bird ID as a random effect.

| | Estimate | Std. Error | z value | Pr(>\|z\|) |
|---|---|---|---|---|
| **Intercept** | -7.94 | 0.96 | -8.27 | <2e-16 *** |
| ***Ad libitum*** | 1.77 | 0.86 | 2.04 | 0.04 * |
| **Food-deprived** | 0.79 | 0.88 | 0.90 | 0.37 |
| **High Protein** | 1.94 | 0.86 | 2.26 | 0.02 * |
| **Low Protein** | 1.37 | 0.86 | 1.60 | 0.11 |
| **trial 3** | 0.008 | 0.006 | 1.36 | 0.17 |

**Table 20S.** Beak wiping reaction to the bait (oat) without control was tested using a generalised linear mixed-effects models (GLMM) with a log link and Poisson distribution, fit by maximum likelihood (Laplace approximation) to test for differences in beak wiping events per minute.The interaction between diet (low protein, high protein), the amount of melanin, and trial were set as fixed effects, while the beak wiping frequency was set as response variable and bird ID as a random effect

| | Estimate | Std. Error | z value | Pr(>\|z\|) |
|---|---|---|---|---|
| **Intercept** | -6.48 | 0.57 | -11.37 | <2e-16 *** |

| | | | | |
|---|---|---|---|---|
| **High Protein-Low Protein** | 0.29 | 0.31 | -0.94 | 0.35 |
| ***Ad libitum*-Food depriv** | -0.49 | 0.33 | -1.48 | 0.14 |
| **Natural Diet-Artificial Diet** | 0.19 | 0.22 | -0.86 | 0.39 |
| **trial 3** | 0.008 | 0.006 | 1.36 | 0.175 |

#### 7. Female fecundity

High- and low-protein diets were repeated in the summer of the following year using families taken from the same Estonian stock. Larvae were maintained as above. In the separate follow-up experiment, a total of 58 females from 17 families, 26 reared on low-protein diet and 31 on high-protein diet, were paired with a randomly chosen non-sibling male from the laboratory stock and allowed to mate and lay eggs. The number of eggs and resulting hatched larvae from each pair was counted. Mating success was measured as if a female produced at least one fertile egg.

Females raised on high-protein diets produced more larvae than those reared on low-protein diets ($t=2.25$, $p= 0.03$; Supplementary Figure 2). The number of eggs laid also showed a non-significant trend to be higher in high-protein females ($t=2.01$, $p= 0.052$). The best fitting model for both measures included female weight, although it did not significantly affect the number of eggs ($t=1.365$, $p = 0.18$) or larvae ($t= 1.00$, $p = 0.32$) produced, and female family as a random factor. Crucially, the rate of failed matings did not differ between the two groups ($z=0.25$, $p=0.80$), suggesting that this pattern was being driven by female egg production, and not mating success.
